# Two TAL effectors of *Xanthomonas citri* pv. *malvacearum* target *GhSWEET15* as the susceptibility genes for bacterial blight of cotton

**DOI:** 10.1101/2024.08.06.606744

**Authors:** Syed Mashab Ali Shah, Fazal Haq, Kunxuan Huang, Qi Wang, Linlin Liu, Ying Li, Yong Wang, Asaf Khan, Ruihuan Yang, Moein Khojasteh, Xiameng Xu, Zhengyin Xu, Gongyou Chen

## Abstract

Bacterial Blight of Cotton (BBC) caused by *Xanthomonas citri* pv. *malvacearum* (*Xcm*) is an important and destructive disease affecting cotton plants. Transcription activator-like effectors (TALEs) released by the pathogen regulate cotton resistance to the susceptibility. In this study, we sequenced the whole genome of *Xcm* Xss-V_2_-18 and identified eight *tal* genes; seven on the plasmids and one on the chromosome. Deletion and complementation experiments of Xss-V_2_-18 *tal* genes demonstrated that Tal1b is required for full virulence on cotton. Transcriptome profiling coupled with TALE-binding element prediction revealed that Tal1b targets *GhSWEET15A04/D04* and *GhSWEET15D02* simultaneously. Expression analysis confirmed the independent inducibility of *GhSWEET15A04/D04* and *GhSWEET15D02* by Tal1b, whereas *GhSWEET15A04/D04* is additionally targeted by Tal1. Moreover, GUS (β*-glucuronidase*) and *Xa10*-mediated HR (hypersensitive response) assays indicated that the EBEs are required for the direct and specific activation of the candidate targets by Tal1 and Ta1b. These findings may advance our understanding of the dynamics between TALEs and EBEs, and decipher a simple and effective DNA-binding mechanism that could lead to the development of more efficient methods for gene editing and transgenic research.

---

Cultivated cotton (*Gossypium* spp.) is one of the most economically important crops, providing a significant source of foodstuff, oil, fiber, and biofuel products worldwide. However, cotton production is constrained by numerous foliar diseases across growing regions. Among these, bacterial blight of cotton (BBC), also called “Angular leaf spot “ caused by *Xanthomonas citri* pv. *malvacearum* (*Xcm*) is a historical and destructive disease. Over the last decade, BBC has emerged as an increasingly important disease in all cotton growing areas around the world (Cox et al., 2017; Jalloul, Sayegh, Champion, & Nicole, 2015; Mijatović, Severns, Kemerait, Walcott, & Kvitko, 2021; Phillips et al., 2017). *Xcm* infects the host plant at any stage of development, from the seedling to the harvesting stage (Elassbli et al., 2021; Mijatović et al., 2021). Typical symptoms include water-soaked lesions that become necrotic, leading to premature defoliation and boll loss. This disease can be destructive even if a single infected plant is observed per 6,000 plants (Jalloul et al., 2015). Therefore, there is a pressing need to understand pathogenicity and host responses to the pathogen in order to develop effective disease prevention and management strategies.

Similar to other Xanthomonads, *Xcm* secretes various effector proteins into plant cells via the type III secretion system (T3SS), acting on various target factors of the host plant, facilitating nutrient acquisition by the pathogen and disease progression (Büttner & Bonas, 2010). Among these effectors, transcription activator-like (TAL) effector (TALE) proteins, that resemble eukaryotic transcription factors, play a crucial role in the complex pathogenesis processes (Bogdanove, Schornack, & Lahaye, 2010). Structurally, TALEs comprise an N-terminal type III secretion signal, a central repeat region (CRR), and C-terminal nuclear localization signals followed by an acidic activation domain. The main differences among these proteins lie in the CRR, which consists of 1.5 to 33.5 nearly identical repeats of 33-34 amino acids. These repeats are conserved except for the 12th and 13th residues in each repeat, known as the repeat variable di-residue (RVD) (Deng et al., 2012). Each RVD corresponds to a specific nucleotide in the promoters of host genes, forming a code where the sequence of RVDs determines the effector-binding element (EBE). Notably, each EBE begins with a nearly invariant thymine, specified by structures immediately N-terminal to the CRR, though the mechanism for this specification is not yet clear (Schreiber & Bonas, 2014).

Following injection, TALE proteins localize in the host cell nucleus, where they recognize and bind to their EBEs in the promoter region of host resistance (*R*) or susceptibility (*S*) genes, inducing their expression (Bogdanove et al., 2010; X. Xu et al., 2022). Based on a handful characterized examples, TALEs exploit host *S* genes, which are crucial for host recognition, subvert plant immunity, and transport nutrients to the pathogen. Most of the known TALE-targeted *S* genes encode transporters and transcriptional factors. For example, three SWEET sugar transporter genes from clade-III are targeted by about ten major TALEs; including *OsSWEET11a*/*Os8N3*/*Xa13* induced by PthXo1 and Tal6b/AvrXa27A at overlapping EBEs (Z. Xu et al., 2023; Yang, Sugio, & White, 2006), *OsSWEET13*/*Os12N3*/*Xa25* targeted by several PthXo2-like TALEs (i.e. PthXo2, PthXo2B/Tal7_PXO61_, PthXo2C/Tal5_LN18_ and Tal7_K74_) (Z. Xu et al., 2019; Zhou et al., 2015), and *OsSWEET14*/*Os11N3*/*Xa41*(*t*) activated by PthXo3, AvrXa7, Tal5/TalF and TalC at overlapping/different EBEs (Antony et al., 2010; Streubel et al., 2013; Tran et al., 2018; Yu et al., 2011). Additionally, TalB targets the rice *S* gene *OsERF#123* in African strains of *Xoo* (Tran et al., 2018). *X. oryzae* pv. *oryzicola* Tal2g targets *OsSULTR3;6*, a sulfate transporter gene important for bacterial leaf streak (Cernadas et al., 2014). Moreover, some TALE-dependent *S* genes are host transcription-factors; for instance, PthXo6 and PthXo7 of *Xoo* induce the expression of rice transcription factors *OsTFX1* and *TFIIA*γ*1*, respectively (Sugio, Yang, Zhu, & White, 2007).

Relatively few TALE-targeted genes have been identified in the interaction of cotton with *Xcm*. Avrb6 was first characterized, which induces the expression of *GhSWEET10* in both A and D genomes of cotton (Cox et al., 2017). This upregulation of *GhSWEET10* leads to the efflux of sucrose from the host cell into the apoplast (L.-Q. Chen et al., 2012; Cox et al., 2017), which can potentially serve as a carbon and energy source for bacterial infection or alter the apoplast water potential, resulting to water-soaked lesion development in plant leaves (Streubel et al., 2013; Zhou et al., 2015). Recently, a new TALE; Tal7b, was characterized in the *Xcm* strain XT7, targeting three genes in cotton genomes and conferring susceptibility (Mormile et al., 2024). Identifying the pathogenicity determinants in the pathogen and its corresponding targets in tetraploid cotton remains a challenge. In our previous study we found that *Xcm* Xss-V_2_-18 encods six *tal* genes based on the southern blotting and their complete sequences were obtained via Tn5 transposon. Reduced water soaking symptoms suggested that Tal2 is a major TALE in Xss-V_2_-18 (Haq et al., 2020). However in this study, whole genome sequencing revealed that this strain harbor additional two *tal* genes namely Tal1b and Tal6b (**Table S3 and S4**), which due to their equal sizes with Tal1 and Tal6, respectively, couldn ‘t be isolated. We complemented TALEs in the *tal*-free (M6) strain previously obtained in our laboratory (Haq et al., 2020), to further identify the TALEs responsible for host plant susceptibility (**Figure 1A and Text S1**). Complementation of TALEs was confirmed by immunoblotting (**Figure 1B**). The *Xcm* Xss-V_2_-18 and mutant strains (M4, M5, M6/EV, M6/Tal1 and M6/Tal1b) were then evaluated for virulence by inoculating TM1 cotton leaves and were photographed at 6 dpi (**Text S1**). Xss-V_2_-18, M4 and M5 produced substantial water-soaked lesions in the inoculation sites; however, M6/EV did not elicit any symptoms, suggesting that Tal1 and/or Tal1b are the major virulence factors. The gain of function experiments demonstrated that the introduction of Tal1b but not Tal1 can restore full virulence in the M6 strain (**Figure 1C**), confirming Tal1b alone is the major virulence factor in Xss-V_2_-18.

**Figure 1.**
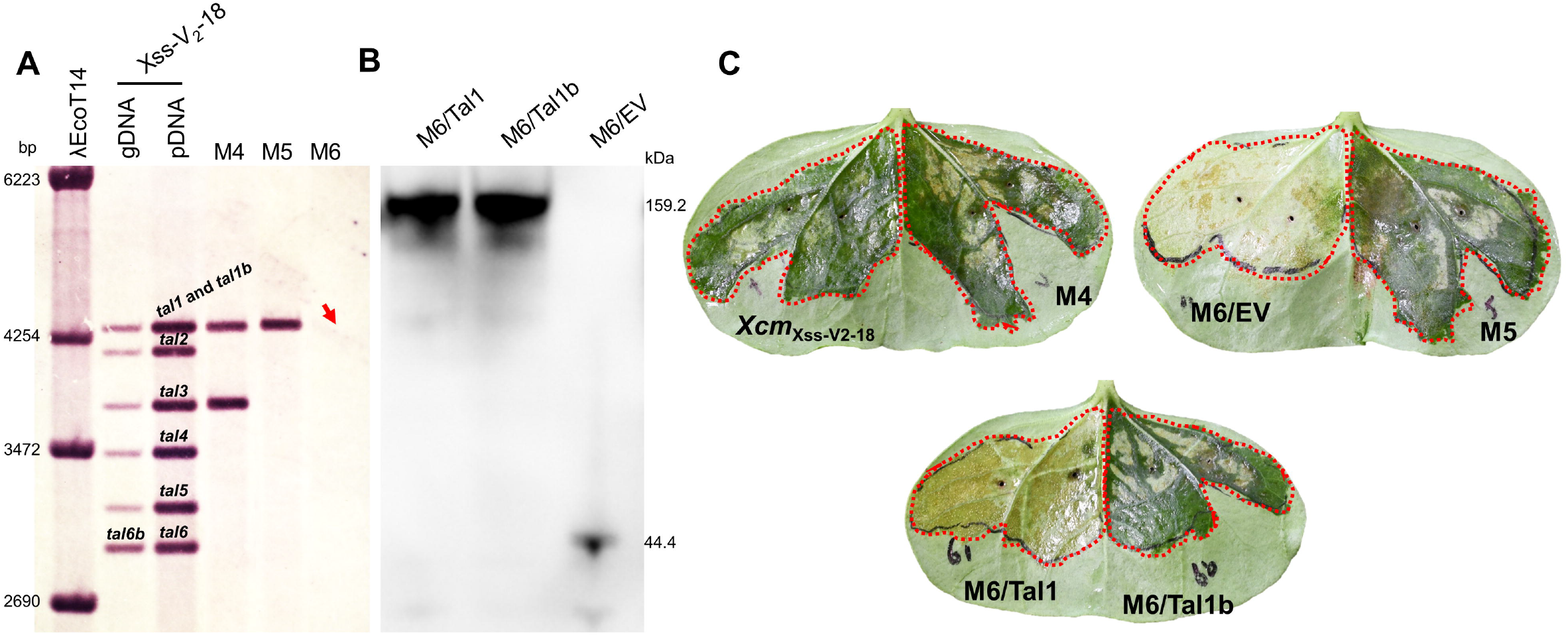
Tal1b is the major virulent TALE in *Xcm* Xss-V_2_-18. **A**) *tal* gene mutants were generated by homologous recombination using the suicide vector pKMS1. The deletion mutants were confirmed by southern blotting (SB) and designated as M4, M5 and M6 (Haq et al., 2020). *tal* genes were named on their order on SB membrane (two *tal* genes of the same size were named “b “). The red arrow indicate deletion of both Tal1 and Tal1b. Strains names are mentioned at the top and the left lanes of the image represent λEcoT14 marker (base pairs; bp). **B**) Tal1 and Tal1b were expressed in M6 mutant, and their expression was confirmed by immunoblotting. Strains are mentioned at the top and marker (kilodalton; kDa) is shown on the right. **C**) *Xcm* and mutant strains were inoculated into TM1 cotton plants, and symptoms were evaluated after 6 dpi. **Abbreviations:** Xss-V_2_-18, wild type strain; M4: Mutant-4; M5: Mutant-5; M6: Mutant-6 (*tal*-free strain); M6/Tal1, M6 having tal1 *in trans*; M6/Tal1b, M6 having tal1b *in trans*; M6/EV, M6 having empty pHZY vector.

To elucidate the underlying mechanism by which Tal1b induces water soaking, we deployed whole genome RNA-sequencing analysis in cotton leaves inoculated with wild-type *Xcm* and the M6 strain (**Figure 2A**). Computational predictions of Tal1 and Tal1b EBEs identified, Tal1b targets *GhSWEET15A04/D04* (Gohir.A04G100600, Gohir.D04G139700) and *GhSWEET15D02* (Gohir.D02G173400) (Z. J. Chen et al., 2020) simultaneously, while *GhSWEET15A04/D04* is also targeted by Tal1 (**Figure 2B**). Concurrently, the mRNA level of *GhSWEET15* (2-ΔΔCt) was determined by qRT-PCR in cotton leaves inoculated with *Xcm*, M4, M5, M6/EV, M6/Tal1 and M6/Tal1b at 24hpi (OD_600_=0.1). The expression of tested *GhSWEET15A04/D04* and *GhSWEET15D02* was induced in *Xcm*, M4 and M5 but not in M6/EV. The M6 carrying Tal1b significantly upregulated *GhSWEET15A04*/*D04* and *GhSWEET15D02* expression to levels comparable to those in *Xcm*, whereas M6 carrying Tal1 induced the expression of *GhSWEET15A04*/*D04* (**Figure 2C**). Importantly, M6 carrying Tal1b, but not other TALEs, restored water-soaked lesions. These results indicate that Tal1b upregulates the expression of *GhSWEET15A04/D04* and *GhSWEET15D02*, mediating water-soaked lesion development.

**Figure 2.**
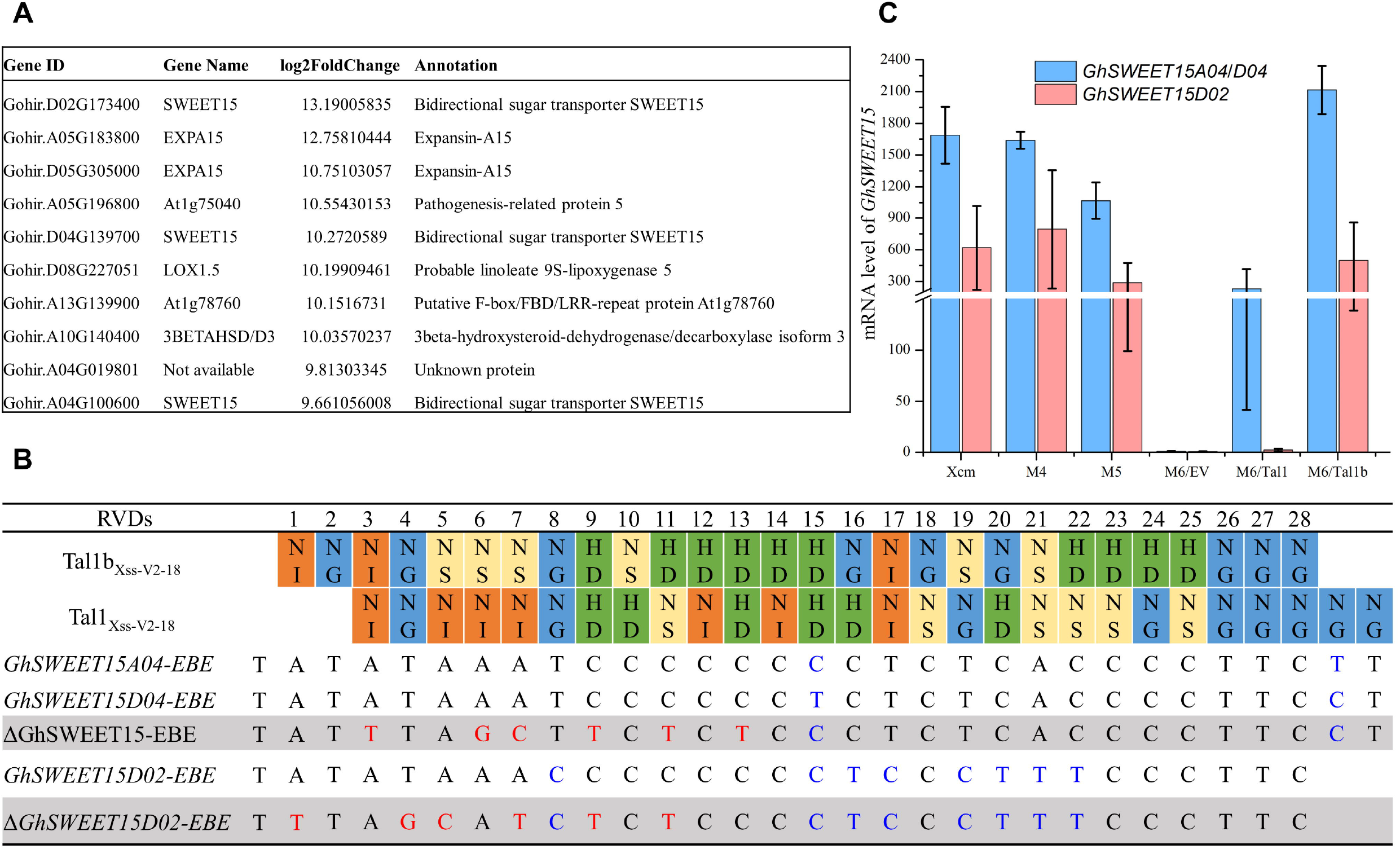
Tal1b and Tal1 targets novel susceptibility genes. **A**) The list of top ten candidate genes from RNA-Sequencing analysis, induced by *Xcm* in comparison with M6, ranked by log2FoldChange. **B**) The RVD sequences of Tal1b and Tal1 and their predicted EBE sites. Colored boxes represent the RVDs of each repeat motif of TALEs, with the number above each box indicating the RVD position within the CRR. ssssThe predicted Tal1b and Tal1 EBE sequences in the promoters of *GhSWEET15A04*/*D04* and *GhSWEET15D02*, identified using TALE-NT, along with the Δ*GhSWEET15* and Δ*GhSWEET15D02* sequences, are listed below the RVDs. Natural SNPs (single nucleotide polymorphisms) and mutated nucleotides are represented in blue and red fonts, respectively. The gray highlighted rows represent the mutated EBEs. **C**) The mRNA level of *GhSWEET15* (2^-ΔΔCt^) was determined by qRT-PCR in cotton leaves transfected with *Xcm*, M4, M5, M6/EV, M6/Tal1b and M6/Tal1 at 24hpi (OD_600_=0.1).

Our working hypothesis was that Tal1 and Tal1b could bind to the predicted EBEs of *GhSWEET15* and activate their expression to benefit the pathogen. Thus, we focused our efforts on studying the interaction of Tal1 and Tal1b with the *GhSWEET115* EBEs. To investigate whether the presence of Tal1 and Tal1b induces the expression of the *GhSWEET15*, we cloned these TALEs into pHB (**Figure 3A**), and their expression in *Nicotiana benthamiana* (*Nb*) was confirmed via western blotting (**Figure 3B**). We also cloned the promoter regions of the *pGhSWEET15A04* (166bp), *pGhSWEET15D04* (169bp), and *pGhSWEET15D02* (70bp) upstream of a *gusA* reporter gene in pCAMBIA1381 (**Figure 3C**). The three promoter::GUS fusions (*pGhSWEET15A04*::GUS, *pGhSWEET15D04*::GUS and *pGhSWEET15D02*::GUS) were individually co-expressed in *Nb* with pHB-Tal1 and pHB-Tal1b through *Agrobacterium* mediated transformation. The effector and reporter plasmids; pHB-PthXo1 and *pOs8N3/OsSWEET11a*::GUS, were used as negative (individually) and positive controls (collectively), respectively. Significant GUS expression was observed when pHB-Tal1 was co-expressed with *pGhSWEET15A04/D04*, and pHB-Tal1b with *pGhSWEET15A04/D04* and *pGhSWEET15D02* (**Figure 3D**). Furthermore, the EBEs of both Tal1 and Tal1b were mutated by substitution, and designated Δ*pGhSWEET15A04/D04* and Δ*pGhSWEET15D02* (**Figure 3E and 3C**). The inoculation of both TALEs with their mutated EBEs failed to induce GUS activity (**Figure 3D**), confirming their binding specificity to the targeted EBEs. These results indicate that both Tal1 and Tal1b have the potential to bind with the *GhSWEET15* EBEs in the promoter region and induce their expression.

**Figure 3.**
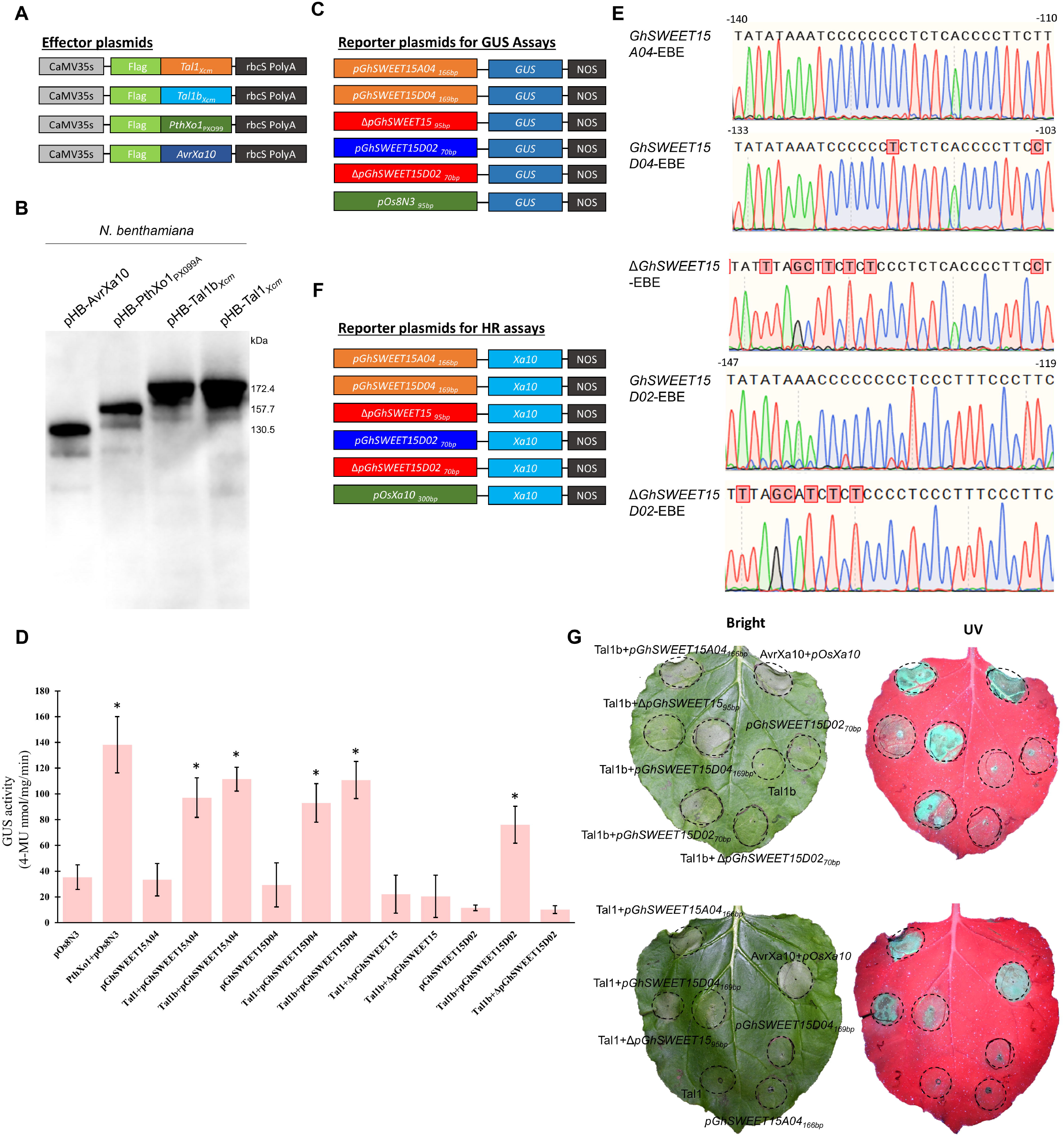
Transactivation assays in *N. benthamiana* confirmed both Tal1 and Tal1b simultaneously target *GhSWEET15* in both A and D genomes. **A**) Schematic representation of the effector constructs contained FLAG-tagged Tal1, Tal1b, PthXo1, and AvrXa10 in pHB vector under the control of the CaMV 35S promoter. **B**) Tal1 and Tal1b protein expression in *N. benthamiana*. Leaves were infiltrated with *Agrobacterium* carrying 35S::FLAG-TALE. Inoculated tissues were collected at 48 hpi and subjected to immunoblotting. **C**) Schematic illustration of reporter constructs containing the *gusA* reporter cassettes driven by the candidate gene promoters cloned in pCAMBIA1381. **D**) GUS activity in *Nb*. Co-inoculation revealed *Xcm* Tal1 and Tal1b can bind to the promoter region upstream of the 5`UTR (untranslated region) of *GhSWEET15A04/D04*, while Tal1b additionally binds to *GhSWEET15D02*, activating GUS activity. The GUS activity was significantly reduced after mutating their EBEs: Δ*GhSWEET15* and Δ*GhSWEET15D02*. PthXo1 and its cognate target *Os8N3* were used as a positive control. Samples were collected at 48-hpi, and GUS activity was calculated. Error bars indicate means ± SD (n = 4) and asterisks indicate significant difference. **E**) Sequencing chromatograms of the *GhSWEET15A04/D04, GhSWEET15D02*, Δ*GhSWEET15*, and Δ*GhSWEET15D02* EBEs are shown below the nucleotide sequences. Red boxes denote mutated nucleotides. Numbers on the sides indicate the EBE proximity to the translational start site. **F**) Schematic diagram of the reporter constructs contained an *Xa10* reporter cassette driven by candidate gene promoters as illustrated. **G**) HR induction in *Nb*. Leaves were infiltrated with *Agrobacterium* containing different combinations of reporter/effector constructs. HR induction was observed and photographed 48 hrs after post infiltration. The circles indicate infiltrated regions of tobacco leaves. These results indicate that Tal1 can bind to the promoter of *GhSWEET15A04/D04*, while Tal1b binds to *GhSWEET15A04/D04* and *GhSWEET15D02*, activating the HR. Additionally, the co-inoculation of Tal1 and Tal1b with the mutated Δ*pGhSWEET15* and Δ*pGhSWEET15D02* did not elicit HR, validating the accuracy of the predicted EBE. The top leaves indicate Tal1b, while the lower leaves show the Tal1 interactions. **Abbreviations**: ribulose-1,5bisphosphate carboxylase, small subunit; polyA, polyadenylation site; NOS, NOS terminator site.

A novel *in vivo* reporter system was developed to investigate further the binding of TALEs-EBEs (**Text S1**), which can trigger an hypersensitive response (HR) once activated *in planta*. In this experiment, *gusA* was replaced with the coding sequence (CDS) of *Xa10*, an executor *R* (resistance) gene in rice that trap *Xoo*-AvrXa10 and confers immunity (Tian et al., 2014; Wang et al., 2017; Zeng et al., 2015), and expression was driven by the *pGhSWEET15A04, pGhSWEET15D04*, and *pGhSWEET15D02* promoters (**Figure 3F**). The constructs pHB-AvrXa10 and *pOsXa10-Xa10* served as positive control (Haq et al., 2021); as *Xa10* encodes a protein that elicits an HR in *Nb*. The reporter constructs were transiently co-expressed in *Nb* with the corresponding effectors. In these screening assays, an HR would develop if the effectors bind to the corresponding EBE in the promoter region, leading to the expression of *Xa10*. When Tal1b was co-expressed with *pGhSWEET15A04*::*Xa10, pGhSWEET15D04*::*Xa10* and *pGhSWEET15D02*::*Xa10*, and Tal1 with *pGhSWEET15A04*::*Xa10* and *pGhSWEET15D04*::*Xa10*, HR-like cell death was apparent starting at 36 hours (**Figure 3G**). To further confirm that *Xa10* activation by Tal1b and Tal1 mediates the HR in *Nb*, we mutated the EBEs (Δ*pGhSWEET15A04/D04* and Δ*pGhSWEET15D02*) (**Figure 3E and 3F**). Leaves inoculated with Tal1/Tal1b+Δ*pGhSWEET15A04/D04* and Tal1b+Δ*pGhSWEET15D02* showed no obvious HR compared to Tal1/Tal1b+*pGhSWEET15A04/D04* and Tal1b+*pGhSWEET15D02* at 2 dpi (**Figure 3G**), confirming the accuracy of predicted EBEs. Collectively, these results provide further evidence that Tal1b binds with all the three promoters *pGhSWEET15A04/D04/D02* and Tal1 binds with *pGhSWEET15A04/D04* via effector binding sites.

In summary, here we identified two novel major TALEs in *Xcm* Xss-V_2_-18; Tal1 and Tal1b, that specifically activate the expression of *GhSWEET15A04/D04* and *GhSWEET15A04/D04/D02*, respectively. We confirmed their binding on overlapping EBEs using different approaches, including GUS reporter assays and *Xa10*-mediated HR assays. TALEs are believed to bind to specific nucleotides; however, in this study, we noticed that Tal1 and Tal1b mediate their interactions through different nucleotides deviating from the currently accepted TALE-Code as Tal6b/AvrXa27b binds to *OsSWEET11a* (Z. Xu et al., 2023). This may be due to selection pressure as a result of evolutionary arm race during plant-pathogen interactions, that forces pathogen to evolve different TALE-codes to confer susceptibility. Tal1 and Tal1b contain 27.5 central repeat units, identical to Tal5 and Tal7b in TX4 and TX7 from Texas, and Tal8 and Tal4 in MSCT1 from Mississippi, respectively (Mormile et al., 2024). The Tal7b of TX4 targets *GhSWEET14a, GhSWEET14b*, and pectin lyase, contributing to symptom development. The *GhSWEET14a* corresponds to *GhSWEET15A04/D04*, while *GhSWEET14b* corresponds to *GhSWEET15D02* (Z. J. Chen et al., 2020). Surprisingly, on the contrary in a recent study, *GhSWEET14a* was not induced in response to Tal5_TX4_ (Mormile et al., 2024). Moreover, we could not detect induction of pectin lyase in our transcriptomic data. These differences in the induction of transcript levels might be due to difference in the strains background. We noticed that in addition to Tal5 of TX4, Tal1 also shares identical RVDs to Tal5 of TX9 which is characterized as less virulent strain (Mormile et al., 2024). Furthermore, Tal1 can induce *GhSWEET15A04/D04*, but could not contribute to the development of water soaking symptoms, may suggest that the evolutionary pressure might be shifting the pathogenicity from minor to robust virulence over time. Despite their geographical distribution, these TALEs are identical across most of the reported pathogenic *Xcm* strains. Collectively, we showed that, Tal1 and Tal1b are major TALEs in *Xcm* Xss-V_2_-18, that can be used for resistance development in cotton by taking advantage of promoter trap strategy.

## Supporting information

Supplementary Informations

## Acknowledgments

We thank Dr. Weiguo Miao at Hainan University (China) for providing *Xcm* strain Xss-V_2_-18. This work was supported by the National Foreign Expert Program (QN2023134007) by Ministry of Science and Technology of the People ‘s Republic of China, the National Natural Science Foundation of China (32361143515), and the Morning Star Postdoctoral Incentive Program by Shanghai Jiao Tong University.

## Data Availability Statement

The sequences of *GhSWEET15A04, GhSWEET15D04* and *GhSWEET15D02* are available in the Phytozome (https://phytozome-next.jgi.doe.gov/info/Ghirsutum_v3_1) with accession number Gohir.A04G100600, Gohir.D04G139700 and Gohir.D02G173400.

## Supporting Information legends

**Text_S1**. Materials and methods

**Figure S1**. Isolation of *tal* genes from *Xcm* strain Xss-V_2_-18. Representative image of colony *in-situ* hybridization results using the *Sph*I fragment from pthXo1 as a probe. The blue lines indicate hybridizing colonies that contain *tal* genes.

**Supplementary Table S1**. Bacterial strains and plasmids used in this study.

**Supplementary Table S2**. Primers used in this study.

**Supplementary Table S3**. Statistical summary of whole-genome sequencing of *Xcm* strain XSS-V_2_-18.

**Supplementary Table S4**. TALE RVD sequences of *Xcm* strain Xss-V_2_-18.

