## Supplementary Informations for "Two TAL effectors of *Xanthomonas citri* pv. *malvacearum* target *GhSWEET15* as the susceptibility genes for bacterial blight of cotton"

^1§^ Former Master degree student, currently working in Shanghai Customs District, Shanghai, China.

^3^ Center for Viticulture and Enology, School of Agriculture and Biology, Shanghai Jiao Tong University, Shanghai, China.

**Materials and methods**

**Plant materials, bacterial strains, and growth conditions**

*Gossypium hirsutum* germplasm TM1 was cultivated in 10-cm circular pots containing Moss peat soil (Pindstrup horticulture Co. Ltd, Shanghai) in a greenhouse at 25°C, 50-65% relative humidity with a 12-hr light/12-hr dark photoperiod. *Nicotiana benthamiana* were cultivated in 8-cm square pots containing Moss peat soil (Pindstrup horticulture Co. Ltd, Shanghai) in a growth chamber at 22°C, 40-50% humidity and with a 12-hr light/12-hr dark photoperiod. Cotton plants inoculated with *Xanthomonas citri* pv. *malvacearum* (*Xcm*) were transferred into a growth chamber at 28°C, 80% humidity, and 280 μmol light with a 16-hr light/8-hr dark photoperiod.

The bacterial strains and plasmids used in this study are listed in **Table S1**. *Xcm* strains were grown in nutrient broth or NB with 1.5% agar (NA) at 28℃. *Escherichia coli* was grown in Luria-Bertani (LB) medium at 37℃, while *Agrobacterium tumefaciens* GV3101 was grown in LB containing rifampicin at 28℃. Antibiotics were used at the following concentrations (μg/mL) when required: ampicillin (Ap) 100, spectinomycin (Sp) 40, rifampicin (Rif) 75, and kanamycin (Km) 25.

**DNA manipulation and plasmid extraction**

Genomic DNA was extracted from a 24-hour culture using the Hipure Bacterial DNA-Kit (Magen, Guangzhou, China). Routine plasmids were isolated with the Plasmid DNA Mini-Kit (GBS Biotechnology, China). DNA gel extraction was performed using miniprep-kit from Generay Biotechnology (Shanghai, China). DNA polymerases, restriction endonucleases, and molecular weight markers were sourced from TaKaRa (Dalian, China). DNA ligase was purchased from Thermo Fisher Scientific (USA). Kits for reverse transcription PCR and fluorescent quantitative PCR were obtained from TransGen Biotech (Beijing, China). Primers were synthesized by Generay (Shanghai, China), and constructs were confirmed via Sanger sequencing by Biosune (Shanghai, China). All kits were used according to the manufacturer’s instructions.

**TALEs mutagenesis, isolations, and expression**

Deletion mutants of *tal* genes were generated via homologous recombination using the suicide vector pKMS1 and confirmed by southern blotting described in our previous study (Haq et al., 2020). For *tal* genes isolation (**Figure S1**), genomic DNA from Xss-V_2_-18 was extracted and digested with *Bam*HI, *Eco*RI, and *Apa*LI. The digested DNA was separated on agarose gel, and the gel slices containing the desired DNA fragments were excised and purified. These fragments were cloned into a *Bam*HI-digested and CIAP-treated pB vector. The recombinant vectors were introduced into dh5α cells. Colonies containing pB-TALE were identified via *in-situ* hybridization using pB-PthXo1_PX099_ *Sph*I fragment as a probe (**Figure S1**). Positive colonies were further validated by *Sph*I restriction digestion and Sanger sequencing using TAL effector primers (**Table S2**). To express *tal* genes in *tal*-free (M6) strain, pB clones with *tal* genes were digested at *Sph*I site to release the CRR and ligated into *Sph*I-linearized, CIP-treated pZY/pZW vectors. The ligated products were transferred into dh5α cells, and the orientation of *tal* genes was analyzed by PCR and confirmed by sequencing. For expression in *Xanthomonas*, pZY vectors containing *tal* genes [pZY-tal1 and pZY-tal1b] were ligated into pHMI at the *Hind*III site and transferred into the M6 strain, resulting in constructs designated as pHZY-tal1 and pHZY-tal1b (**Table S1**).

**Immunoblotting of TALEs in *Xanthomonas* and *N. benthamiana***

To confirm *tal* gene expression in *Xanthomonas*, bacterial cells containing the C-terminally FLAG-tagged *tal* genes (pHZY-tal) were harvested and washed twice with phosphate-buffered saline (PBS). The harvested cells were diluted in PBS and boiled for 8-10 min in water. Proteins were separated in 4-20% gradient gels and transferred to polyvinylidene difluoride (PVDF) membrane using eBLot L2 (Genscript). 5% skimmed milk in TBST [3M NaCl, 1M Tris-HCl (PH=8.0), 0.5% tween] was used to block the membrane for 1 hour shaking at room temperature. TALEs expression were detected by immunoblotting using mouse anti-flag as the primary antibody, followed by goat anti-mouse IgG (H + L). Protein bands were visualized using EasySee Western Kit (Transgene) on Clinx (science instruments co. ltd).

In *Nb*, infiltrated leaf discs were collected at 48 hpi and macerated in native extraction buffer [50 mM Tris-MES, 1 mM MgCl2, 0.5 M sucrose and 10 mM EDTA pH 8.0] plus freshly added 5 mM DTT and protease inhibitor cocktail. The proteins were separated and detected as described above.

**Identification of virulent TALEs and their cognate targets**

The major virulent TALEs were identified by *tal* genes deletion, isolation (**Figure S1**), cloning, and complementation, following the previously described procedure (Haq et al., 2020; Shah et al., 2019). Pathogenicity assays were done by inoculating bacterial suspensions OD_600_ = 0.1 in two weeks old cotton cultivar TM-1, as described in our previous study (Haq et al., 2020). Leaf phenotypes were examined until 6-days after inoculation. TALEs targets were identified by RNA-Seq analysis (Shah et al., 2023) in cotton leaves transfected with wild-type *Xcm* and the M6 mutant strain [three independent biological replicates]. Computational prediction of major TALEs EBE were made on genes upregulated in *Xcm* compared to M6, using TALE-NT 2.0 Target Finder tool. The EBE sequences were confirmed by sequencing the promoter region of TM1 cotton plants. Expression analysis was done via qRT-PCR to determine the independent inducibility of the candidate genes by the major TALEs.

**Transactivation assays in *N. benthamiana***

For GUS (*β-glucuronidase*) and *Xa10*-HR assays in *N. benthamiana*, *Xcm* Tal1 and Tal1b were cloned at *Bam*HI site in binary vector pHB, which contains the 35S promoter and 10XFlag epitope upstream of the polylinker (**Figure 3A**). TALEs protein expression in *N. benthamiana* was detected by western blotting (**Figure 3B**). For reporter constructs, short DNA fragments (~169bp) upstream of the translation start codons of *OsSWEET11*, *GhSWEET15A04,* and *GhSWEET15D04*, containing EBEs recognized by TALEs, were PCR amplified and fused with *gusA* and *Xa10* in pCAMBIA1381 vector (**Figure 3C and 3F**). For cloning the promoter regions of *GhSWEETD02*, Δ*GhSWEET15* (Δ; having mutated EBEs), and Δ*GhSWEETD02* (Δ; having mutated EBEs), long-oligos were annealed and ligated in pCAMBIA1381 vector to generate the reporter constructs (**Figure 3C and 3F**). Primers used for these constructions are provided in **Table S2**. *A. tumefaciens* transformants with effector and reporter constructs were mixed at a 1:1 ratio (OD_600_ = 1.0) and infiltrated into 5- to 7-week-old *N. benthamiana* leaves using needleless syringe. For GUS assays, four leaf discs (1 cm) were collected at 2 dpi, and GUS activity was measured using 4-methylumbelliferyl- *β* –glucuronide as described by Haq et al. (2021). For *Xa10*-mediated HR assays, symptom development was monitored visually and photographed 48 hours after infiltration.

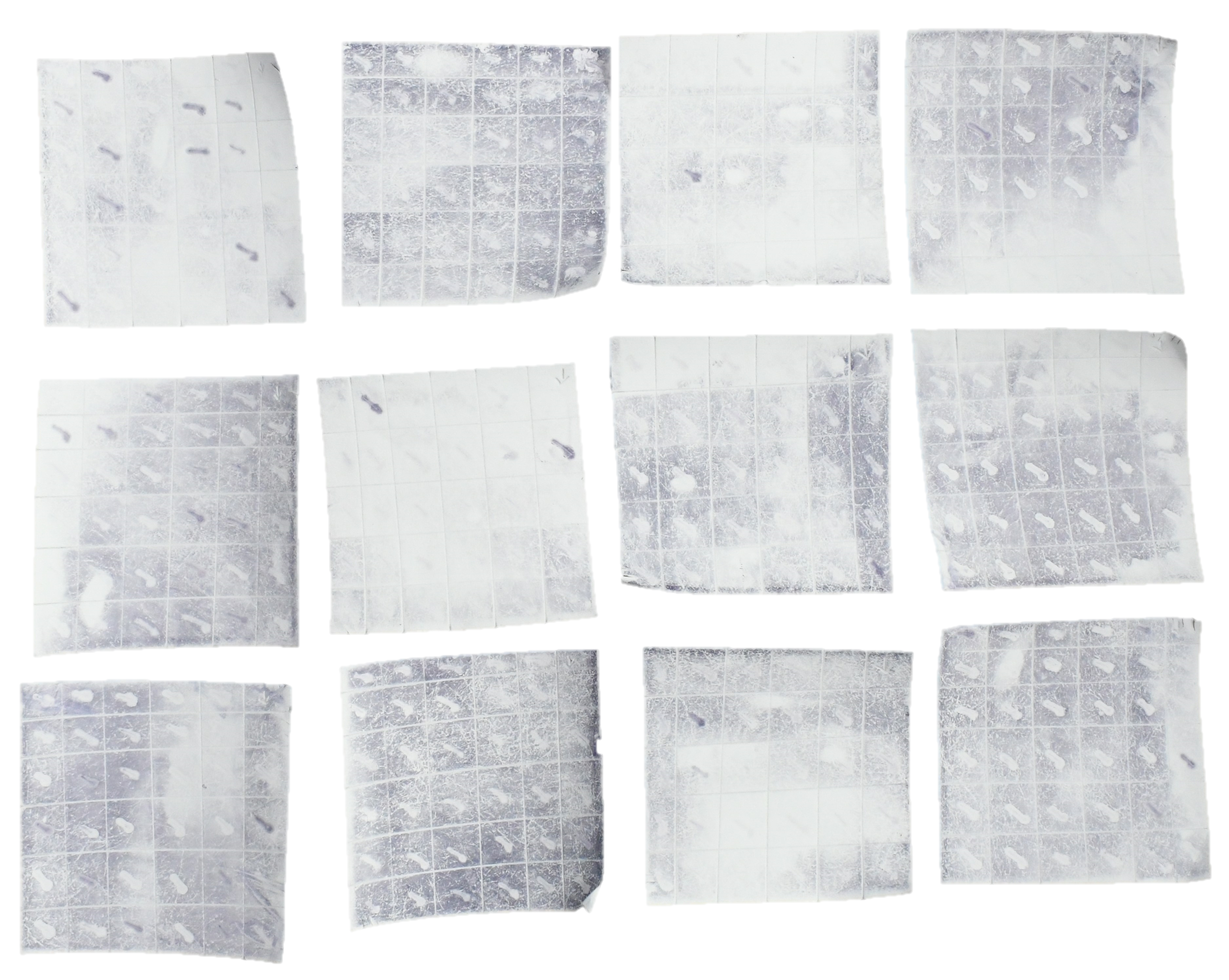

**Figure S1.** Isolation of *tal* genes from *Xcm* strain Xss-V_2_-18. Representative image of colony *in-situ* hybridization results using the *Sph*I fragment from pthXo1 as a probe. The blue lines indicate hybridizing colonies that contain *tal* genes.

**Supplementary Table S1. Bacterial strains and plasmids used in this study.**

| **Strains and Plasmids** | **Features** | **Source** |
| --- | --- | --- |
| **Strains** |  |  |
| ***Xanthomonas citri* pv. *malvacearum*** | |  |
| Xss-V_2_-18 | *Xcm* wild type strain, causes bacterial blight of cotton, from Hainan, China | This Lab |
| M4 | M4 is derived from Xss-V_2_-18 with a deletion of at least five *tal* genes | This Lab |
| M5 | M5 is derived from Xss-V_2_-18 with a deletion of at least six *tal* genes | This Lab |
| M6 | M6 is *tal* free strain derived from Xss-V_2_-18 | This Lab |
| M6/Ev | M6 strain having pHZY empty vector *in trans,* Ap^r^ SP^r^ | This study |
| M6/Tal1 | M6 strain having Tal1 (pHZY-tal1) *in trans*, Ap^r^ SP^r^ | This study |
| M6/Tal1b | M6 strain having Tal1b (pHZY-tal1b) *in trans*, Ap^r^ Sp^r^ | This study |
| ***Escherichia coli*** |  |  |
| DH5α | F- *endA1, thi-1, recA1,* Φ80lacZ, ΔM15 | Clontech |
| ***Agrobacterium tumefaciens*** | |  |
| GV3101 | C58, pTic58DT-DNA, Rif^r^ | This Lab |
| **Plasmids** |  |  |
| pB | Ap^r^, pBluescript II phagemid, pUC derivative | This Lab |
| pKMS1 | Km^r^, pUC18 polylinker, *mob, oriV, sacB* | This Lab |
| pHM1 | Sp^r^, cosmid vector, *cos, parA, IncW* | This Lab |
| pZY/pZW | Ap^r^, pBluescript II with FLAG-tag N- (831-bp) and C- (363-bp) terminus of avrXa10. Lacks the *Sph*I fragment containing the CRR; for expression of TAL effector genes | This Lab |
| pB-tal1 | Ap^r^, pB containing tal1 at *Bam*HI site | This study |
| pB-tal1b | Ap^r^, pB containing tal1b at *Bam*HI site | This study |
| pZY-tal1 | Ap^r^, CRR of tal1 from *Sph*I digested pB-tal1 were ligated into *Sph*I-linearized, CIP-treated pZY vector | This study |
| pZY-tal1b | Ap^r^, CRR of tal1b from *Sph*I digested pB-tal1b were ligated into *Sph*I-linearized, CIP-treated pZY vector | This study |
| pHZY-tal1 | Ap^r^ Sp^r^, pZY with tal1 in pHM1 at *Hin*dIII site | This study |
| pHZY-tal1b | Ap^r^ Sp^r^, pZY with tal1b in pHM1 at *Hin*dIII site | This study |
| pHB | Binary vector, double 35S promoter, 10X N-terminal FLAG tag, Km^r^ | This Lab |
| pHB-AvrXa10 | AvrXa10 cloned in frame with N-terminal flag-tag in pHB, Km^r^ | This Lab |
| pHB-PthXo1 | PthXo1 cloned in frame with N-terminal flag-tag in pHB, Km^r^ | This Lab |
| pHB-Tal1 | Tal1 cloned in frame with N-terminal flag-tag in pHB, Km^r^ | This study |
| pHB-Tal1b | Tal1b cloned in frame with N-terminal flag-tag in pHB, Km^r^ | This study |
| pCAMBIA1381-*gusA* | Binary vector for transient expression, Km^r^ | This Lab |
| pGhSWEET15A04 | 166 bp promoter region of *GhSWEET15A04* (Gohir.A04G100600) cloned upstream of *gusA* in pCAMBIA1381 at *Bam*HI and *Pst*I sites | This study |
| pGhSWEET15D04 | 169 bp promoter region of *GhSWEET15D04* (Gohir.D04G139700) cloned upstream of *gusA* in pCAMBIA1381 at *Bam*HI and *Pst*I sites | This study |
| pGhSWEET15D02 | 70 bp promoter region of *GhSWEET15D02* (Gohir.D02G173400) cloned upstream of *gusA* in pCAMBIA1381 at *Bam*HI and *Pst*I sites | This study |
| pΔGhSWEET15 | 95 bp promoter fragment [Annealed Oligos] of *GhSWEET15A04/D04* having mutation in the predicted EBEs cloned upstream of *gusA* in pCAMBIA1381 at *Bam*HI and *Pst*I sites | This study |
| pΔGhSWEET15D02 | 70 bp promoter fragment [Annealed Oligos] of *GhSWEET15D02* having mutation in the predicted EBEs cloned upstream of *gusA* in pCAMBIA1381 at *Bam*HI and *Pst*I sites | This study |
| pOs8N3 | *gusA* CDS in pCAMBIA1381, driven by *Os8N3* promoter, Km^r^ | This study |
| pGhSWEET15A04-Xa10 | *Xa10* CDS in pCAMBIA1381, driven by *GhSWEET15A04* promoter, Km^r^ | This study |
| pGhSWEET15D04-Xa10 | *Xa10* CDS in pCAMBIA1381, driven by *GhSWEET15D04* promoter, Km^r^ | This study |
| pGhSWEET15D02-Xa10 | *Xa10* CDS in pCAMBIA1381, driven by *GhSWEET15D02* promoter, Km^r^ | This study |
| pΔGhSWEET15-Xa10 | *Xa10* CDS in pCAMBIA1381, driven by Δ*GhSWEET15A04/D04* promoter, Km^r^ | This study |
| pΔGhSWEET15D02-Xa10 | *Xa10* CDS in pCAMBIA1381, driven by Δ*GhSWEET15D02* promoter, Km^r^ | This study |
| pOsXa10-Xa10 | *Xa10* CDS in pCAMBIA1381, driven by *OsXa10* promoter, Km^r^ | This Lab |

**Abbreviations:** Ap^r^, ampicillin resistance; Km^r^, kanamycin resistance; Sp^r^, spectinomycin resistance; Rif^r^, Rifampicin resistance.

**Supplementary Table S2. Primers used in this study.**

**General Primers**

| **Primer Name** | **Sequence 5' to 3'** | **Description** |
| --- | --- | --- |
| RV-M | AGCGGATAACAATTTCACACAGG | Universal primers used for the confirmation and sequencing of the construct |
| M13-47 | CGCCAGGGTTTTCCCAGTCACGAC |  |
| pHB-R | GCATTGAACTTGACGAACGTTGTCGA | Amplifying ~900bp fragment along with Tal-C-F (primer), used in screening positive colonies of pHB having TALE *Bam*HI fragment |
| GUS-R | CGCGATCCAGACTGAATG | Used for screening positive colonies carrying the cloned fragment in pCAMBIA1381-*gusA* vector. |
| Xa10-R | TCAGACGGGGGAAATCTCCT | Used for screening positive colonies carrying the cloned fragment in pCAMBIA1381-*Xa10* vector |

**Promoter Cloning Primers**

| **Primer Name** | **Sequence 5' to 3'** | **Description** |
| --- | --- | --- |
| Os11R2-PstI | AACTGCAGCCATGGCTCAGTGTTTATATAG | Used for amplifying the 95 bp promoter region of rice *OsSWEET11* (*Os8N3*) flanking the EBE for GUS assays |
| Os11F3-EcoRI | CCGGAATTCCACAAGAAAAAAAAGCAAAGGT |  |
| AD04sp-BamHI-F | CGCGGATCCTGTGCTAAGCGTTTGCACTTG | Used for amplifying the 166 bp promoter fragment of *GhSWEET15A04* and *GhSWEET15D04* flanking the EBEs and cloned in pCAMBIA1381 at *Bam*HI and *Pst*I sites |
| AD04sp-PstI-R | AACTGCAGGCTTTTGCTCATGAGGGGTTG |  |

**Note:** Nucleotides denoted with red fonts indicate restriction enzyme sites.

**Long Oligos of Promoter Sequences**

| **Primer Name** | **Sequence 5' to 3'** | **Description** |
| --- | --- | --- |
| SW15D02-Oligos-F | GATCCccacgccaccattcctccctatatataaacccccccctccctttcccttcccctcagctcacccacatcaCTGCA | Used for annealing and ligating the 70 bp promoter fragment of *GhSWEET15D02* in pCAMBIA1381 at *Bam*HI and *Pst*I sites |
| SW15D02-Oligos-R | GtgatgtgggtgagctgaggggaagggaaagggaggggggggtttatatatagggaggaatggtggcgtggG |  |
| SW15D02-MOligos-F | GATCCccacgccaccattcctccctatTtaGCaTcTcTcccctccctttcccttcccctcagctcacccacatcaCTGCA | Used for annealing and ligating the 70 bp promoter fragment of *GhSWEET15D02* with mutated EBE in pCAMBIA1381 at *Bam*HI and *Pst*I sites |
| SW15D02-MOligos-R | GtgatgtgggtgagctgaggggaagggaaagggaggggAgAgAtGCtaAatagggaggaatggtggcgtggG |  |
| GhSWEET15-M.Oligos-F | GATCCagaatgaagcttacatcaccaatttaccatcctccaccttttcccctATtatTtaGCtTcTcTccctctcaccccttcCtttcaacccctcatgaCTGCA | Used for annealing and ligating the 95 bp promoter fragment of *GhSWEET15A04/D04* with mutated EBE in pCAMBIA1381 at *Bam*HI and *Pst*I sites |
| GhSWEET15-M.Oligos-R | GtcatgaggggttgaaaGgaaggggtgagagggAgAgAaGCtaAataATaggggaaaaggtggaggatggtaaattggtgatgtaagcttcattctG |  |

**Note:** Nucleotides in red indicate restriction enzyme sites. Nucleotides in capital black letters represent substitutions.

**TAL-Effectors Sequencing Primers**

| **Primer Name** | **Sequence 5' to 3'** | **Description** |
| --- | --- | --- |
| TALN18-XF | AGAGGCGACACACGAAGCGA | Used to sequence the TALE genes CRR |
| TALN18-XR | CACTGCATCCAGCGCAGGAC |  |
| TALN18-R | TCGCTTCGTGTGTCGCCTCT | Used to sequence the N-terminus of TALE genes |
| Tal-C-F | AGCATTGTTGCCCAGTTAT | Used to sequence the C-terminus of TALE genes |

**qRT-PCR primers**

| **Primer Name** | **Sequence 5' to 3'** | **Description** |
| --- | --- | --- |
| GhUBQ1-F | CTGAATCTTCGCTTTCACGTTATC | Reference gene/Endogenous control for qRT-PCR |
| GhUBQ1-R | GGGATGCAAATCTTCGTGAAAAC |  |
| GhSW15D-F | ACCCTTTCAGCCCCACACTC | For Gh*SWEET15D02* (Gohir.D02G173400) expression analysis via qRT-PCR |
| GhSW15D-R | GCCAAGGAATGGTGATCTGCC |  |
| GhSW15A-F | TGCCTCTGAAGTTCACCCCG | For Gh*SWEET15A04* (Gohir.A04G100600) and *GhSWEET15D04* (Gohir.D04G139700) expression analysis via qRT-PCR |
| GhSW15A-R | CTCCCCTGTGGGTTCGTTGT |  |

**Supplementary Table S3. Statistical summary of whole-genome sequencing of *Xcm* strain XSS-V_2_-18**

The *Xcm* strain Xss-V_2_-18 genome was sequenced using NovaSeq 6000 plus long-read Oxford Nanopore PromethION sequencing platform and assembled de novo (Shah et al., 2021). The characteristics of the assembled genome are presented in the table below.

| **Genome** | **Size(bp)** | ***tal*-gene number** | ***tal*-gene** |
| --- | --- | --- | --- |
| **Chromosome** | 4998219 | 1 | *tal6b* |
| **Plsamid1** | 91181 | 5 | *tal1*, *tal2*, *tal4*, *tal5*, *tal6* |
| **Plasmid2** | 15245 | 0 | none |
| **Plsamid3** | 43949 | 2 | *tal1b*, *tal3* |

**Supplementary Table S4. TALE RVD sequences of *Xcm* strain Xss-V_2_-18**

| **TALE** | **Size(bp)** | **RVDs** | **1 2 3 4 5 6 7 8 9 10 11 12 13 14 15 16 17 18 19 20 21 22 23 24 25 26 27 28** |
| --- | --- | --- | --- |
| Tal1 | 4368 | 27.5 | NI-NG-NI-NI-NI-NG-HD-HD-NS-NI-HD-NI-HD-HD-NI-NS-NG-HD-NS-NG-NS-NG-NS-NG-NG-NG-NG-NG |
| Tal1b | 4362 | 27.5 | NI-NG-NI-NG-NS-NS-NS-NG-HD-NS-HD-HD-HD-HD-HD-NG-NI-NG-NS-NG-NS-HD-HD-HD-HD-NG-NG-NG |
| Tal2 | 4164 | 25.5 | NI-NG-NI-NI-NI-NG-NG-NS-NG-NS-NS-NG-NS-NG-HD-NS-HD-NS-HD-NS-NG-NG-NG-NG-NG-NG |
| Tal3 | 3756 | 21.5 | HD-NI-NG-NI-NI-NS-NG-NG-NI-NG-NS-HD-NS-HD-NS-NG-NS-NG-HD-NG-NG-NG |
| Tal4 | 3450 | 18.5 | HD-NI-NG-NI-NI-NI-HD-HD-HD-NS-NS-HD-HD-NS-NS-NG-NS-NG-NG |
| Tal5 | 3141 | 15.5 | NI-NI-NI-NN-NI-NS-HD-NG-NN-NS-NN-NN-HD-NG-N*-NN |
| Tal6 | 2940 | 13.5 | NI-NI-NI-NN-NG-NS-HD-NG-HD-NS-NG-HD-HD-NG |
| Tal6b | 2934 | 13.5 | NI-NG-NI-HD-NG-NG-NG-NG-HD-NS-HD-HD-NG-NG |
